# Viral manipulation of functionally distinct neurons from mice to humans

**DOI:** 10.1101/808170

**Authors:** Douglas Vormstein-Schneider, Jessica Lin, Kenneth Pelkey, Ramesh Chittajallu, Baolin Guo, Mario Arias Garcia, Kathryn Allaway, Sofia Sakopoulos, Gates Schneider, Olivia Stevenson, Josselyn Vergara, Jitendra Sharma, Qiangge Zhang, Tom Franken, Jared Smith, Leena Ibrahim, Kevin Mastro, Ehsan Sabri, Shuhan Huang, Emilia Favuzzi, Timothy Burbridge, Qing Xu, Lihua Guo, Ian Vogel, Vanessa Sanchez, Giuseppe Saldi, Xiaoqing Yuan, Kareem Zaghloul, Orrin Devinsky, Bernardo Sabatini, Renata Batista-Brito, John Reynolds, Guoping Feng, Zhanyan Fu, Chris McBain, Gord Fishell, Jordane Dimidschstein

## Abstract

Recent success in identifying gene regulatory elements in the context of recombinant adeno-associated virus vectors have enabled cell type-restricted gene expression. However, within the cerebral cortex these tools are presently limited to broad classes of neurons. To overcome this limitation, we developed a strategy that led to the identification of multiple novel enhancers to target functionally distinct neuronal subtypes. By investigating the regulatory landscape of the disease gene Scn1a, we identified enhancers that target the breadth of its expression, including two that are selective for parvalbumin and vasoactive intestinal peptide cortical interneurons. Demonstrating the functional utility of these elements, we found that the PV-specific enhancer allowed for the selective targeting and manipulation of these neurons across species, from mice to humans. Finally, we demonstrate that our selection method is generalizable to other genes and characterize four additional PV-specific enhancers with exquisite specificity for distinct regions of the brain. Altogether, these viral tools can be used for cell-type specific circuit manipulation and hold considerable promise for use in therapeutic interventions.

---

Large-scale transcriptomic studies are rapidly revealing where and when genes associated with neuropsychiatric disease are expressed within specific cell types (1–4). Approaches for understanding and treating these disorders will require methods for targeting and manipulating specific neuronal subtypes. Thus, gaining access to these populations in non-human primates and humans has become paramount. AAVs are the method of choice for gene delivery in the nervous system but have a limited genomic payload and are not intrinsically selective for particular neuronal populations (5). We and others have identified short regulatory elements capable of restricting viral expression to broad neuronal classes. In addition, systematic enhancer discovery has been accelerated by the recent development of technologies allowing for transcriptomic and epigenetic studies at single-cell resolution (6–12). Despite these advances, the search space for enhancer selection remains enormous and to date success has been limited. To focus our enhancer selection, we chose to specifically examine the regulatory landscape of *Scn1a*, a gene expressed in distinct neuronal populations and whose disruption is associated with severe epilepsy (13). Combining single-cell ATAC-seq data with sequence conservation across species, we nominated ten candidate regulatory sequences in the vicinity of this gene. By thoroughly investigating each of these elements for their ability to direct viral expression, we identified three enhancers that collectively target the breadth of neuronal populations expressing *Scn1a*. Among these, one particular short regulatory sequence was capable of restricting viral expression to parvalbumin-expressing cortical interneurons (PV cINs). To fully assess the utility of this element beyond reporter expression, we validated it in a variety of contexts, including synaptic tagging, calcium imaging, as well as opto- and chemo-genic approaches, both *ex vivo* and *in vivo*. Moreover, we showed that this element allows for the selective targeting of PV cINs both during development and across species including rodents, non-human primates and humans. Demonstrating that this approach provides a generalizable strategy for enhancer discovery, we further selected twenty-five regulatory elements in the vicinity of seven genes enriched in PV INs. From these we identified an additional four PV-specific regulatory elements, each of which had remarkably selective expression within specific brain regions. Together we demonstrate the utility of a variety of functionally-tested tools that can be immediately utilized across animal models. These viral reagents can be employed to interrogate how functionally distinct neuronal cell-types are affected in the context of neurological, neurodevelopmental and neurodegenerative disease in non-human primates. Ultimately, these may provide the means to therapeutically normalize pathological neuronal activity or gene expression in specific neuronal populations.

## RESULTS

### Identification of Scn1a enhancers

Scn1a encodes for Nav1.1, a sodium channel expressed in three non-overlapping neuronal populations, fast-spiking cortical interneurons expressing parvalbumin (PV cINs), dis-inhibitory cortical interneurons expressing the vaso-intestinal peptide (VIP cINs) and layer 5 pyramidal neurons (14–16). Haploinsufficiency or pathogenic variants of *Scn1a* cause Dravet Syndrome, a common and intractable form of epileptic encephalopathy characterized by the early onset of seizures (17–19). To devise a genetic strategy to target the distinct cortical populations expressing this gene, we developed an integrative method to systematically identify candidate enhancers within this locus. Regulatory sequences were selected based on the following three criteria. First, it has been posited that the proximity of the enhancer to the transcriptional start site (TSS) of a gene scales directly to the level of expression (20). Therefore, to identify enhancers capable of driving functional levels of transgenes, we examined the intergenic and intronic regions of *Scn1a* closest to its TSS. Second, the location of active enhancers within a given cell type correlates with chromatin accessibility (21,22). To assess the chromatin landscape of the cellular populations expressing *Scn1a*, we collected interneurons from the visual cortex, using *Dlx6a^cre^;Sun1-eGFP* transgenic mice. Subsequent to isolating nuclei, we performed single-cell ATAC-seq profiling (23,24) and used the SnapATAC analysis pipeline (see methods) to identify differentially accessible chromatin regions in each of the four major classes of cortical interneurons (Figure 1a-c and supplementary figure 1). Third, as regulatory elements are subject to positive selection pressure, we identified sequences showing the highest conservation across mammalian species, including humans (25–29). Thus, to nominate enhancers with therapeutic potential, we focused on ten selectively accessible intronic and intergenic regions near the TSS of Scn1a that are highly conserved through evolution (E1 to E10 – figure 1d and supplementary table 1).

**Figure 1.**
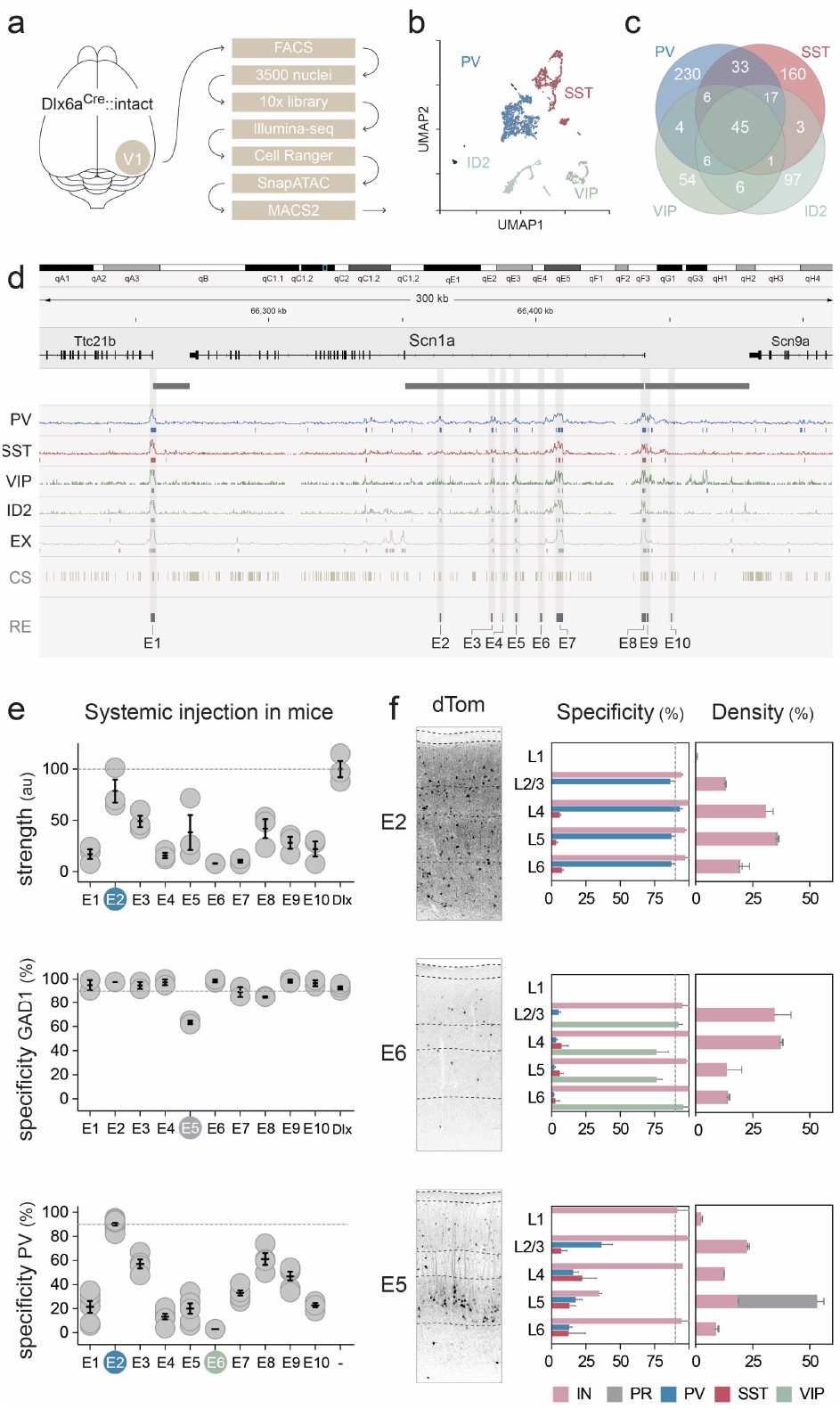
Identification of Scn1a enhancers. (**a**) Schematic representation of the scATAC-seq pipeline. Interneurons were collected from the visual cortex of adult Dlx6a^Cre^::Sun1-eGFP mice. (**b**) Plot of the 3500 nuclei in UMAP space. The clusters obtained from the SnapATAC pipeline were lumped into the four cardinal classes of interneuron. (**c**) Venn diagram showing the number of unique and shared peaks across the four interneuron populations. (**d**) Schematic representation of the enhancer selection method at the Scn1a locus (see methods section for complete description). (**e, f**). Adult mice were injected systemically with the indicated rAAV-E[x]-dTomato and analyzed 3 weeks post-injection. IHC for the reporter and indicated markers in the S1 cortex was used to assess the strength of expression of the reporter (e – upper panel) and the specificity of expression of the viral reporter for the indicated markers (all other panels). Representative fluorescent images of the indicated viral reporter in the somatosensory cortex (f – left panels). Dashed lines represent the limits of anatomical structures. Scale bars represent 100um. On the graphs, the dots represent individual measurements and the lines represent average +/− s.e.m. Values for specificity, sensitivity and strength are listed in the supplementary table 2.

To examine the ability of the candidate enhancers to target the neuronal populations that express Scn1a, each sequence was inserted into an rAAV-backbone containing a minimal promoter upstream of a red fluorescent reporter (rAAV-E[x]-dTomato). From these constructs, rAAVs were then produced with the PHPeB capsid (30) and systemically injected into adult mice. After 3 weeks, all viruses showed strong and sparse expression within the cortex, as well as across multiple brain regions. Except for E5, the vast majority of virally-labelled cells expressed the pan-interneuron marker Gad1. However, the degree of colocalization for PV within cortical neurons varied, ranging from over 90% for E2 to below 5% for E6, with all remaining enhancers displaying intermediate levels of PV specificity (figure 1e and supplementary figures 2). Next, we further examined the identity and layer distribution of the neuronal populations captured by the E2, E5 and E6 enhancers. Consistent with their layer distribution, co-localization analysis with various markers revealed that the E2 regulatory element restricted the expression of the viral reporter to PV cINs, while E6 was selective for VIP interneurons. By contrast, the E5 regulatory element, while sparsely labeling interneurons across all layers, had a notable enrichment for pyramidal neurons in layer 5 (figure 1f). These data indicate that a significant fraction of the cortical expression profile of Scn1a is mirrored by the collective expression of 3 enhancers. These regulatory elements therefore account for largely non-overlapping expression in populations of neurons with distinct functions and developmental origins. The viral tools developed here thus provide a means of dissecting neuronal subtypes and can be used to study their normal function, as well as abnormalities in diseased cortex.

### Viral targeting of PV cINs in mice

With 90% specificity for PV cINs, the E2 regulatory element allows for the targeting of fast-spiking neurons, which constitute 40% of all cortical interneurons. These neurons exert a strong level of inhibition over local networks and their disfunction has been directly implicated in neurological and neuropsychiatric disorders, including Dravet syndrome, focal epilepsy, ASD and schizophrenia (31–35). As such, gaining control over their activity is of particular interest for both fundamental research and clinical applications. Thus, we focused our efforts on characterizing the E2 regulatory element to develop a viral tool with broad utility. Adult mice systemically-injected with rAAV-E2-dTomato showed detectable expression of the viral reporter after one week and reached a high and stable level of expression after 3 weeks. Immunohistochemistry and in situ hybridization consistently showed that ~90% of virally labeled cells were PV INs in the cortex. Conversely, on average 75% of PV cINs expressed the viral reporter, reaching a maximum of 93% (figure 2a and 2b). This suggests that E2 is capable of targeting all PV cINs without bias for layers or subtypes. Consistent with these findings, slice recordings from mice showed that the neurons expressing the viral reporter exhibited electrophysiological properties that are characteristic of fast-spiking PV cINs both within the primary somatosensory cortex (S1) and the pre-frontal cortex (PFC – figure 2b and supplementary figure 3).

**Figure 2.**
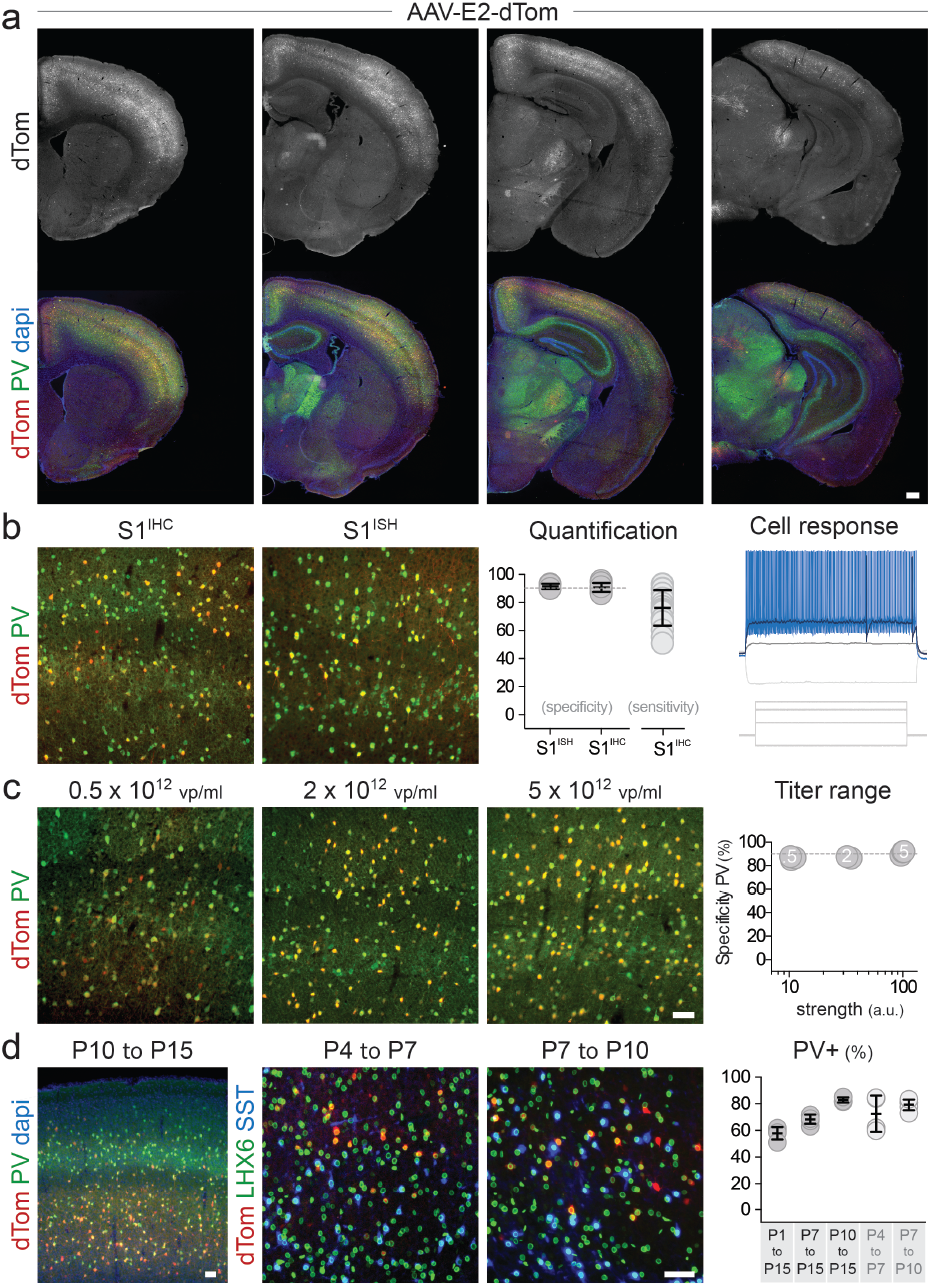
Viral targeting of PV cINs in mice. Adult mice were injected systemically **(a,b)** or locally **(c)** with rAAV-E2-dTomato and analyzed 3 weeks post-injection by IHC or ISH for both the reporter and PV. **(b, right panel)** Representative recording from slice analysis of the intrinsic properties of virally labeled neurons. **(d)** Mice were injected locally with rAAV-E2-dTomato and analyzed at the indicated developmental stages for the reporter and the indicated markers. Scale bars represent 250um **(a)** and 50um **(b,d,e)**. On the graphs, the dots represent individual measurements and the lines represent average +/− s.e.m. Values for specificity, sensitivity and strength are listed in the supplementary table 2.

Although the viral reporter was predominantly confined to the cortex, some positive cells were observed in other brain regions closely corresponding to areas of Scn1a expression. E2 maintained high specificity for PV-expressing neurons within the primary visual cortex (V1) and cingulate cortex, subiculum, hippocampal CA1 and substantia nigra pars reticulata (supplementary figure 4a). Importantly, virtually no viral reporter expression was observed outside of the brain, with the exception of a few cells observed in the liver (which is expected upon systemic delivery of any AAVs) and in the lungs (where Scn1a is expressed at a low level – supplementary figure 4b). This shows that despite systemic delivery, the E2 vector can be used to selectively target PV-expressing neurons in various brain regions, with insignificant off-target expression outside of the central nervous system.

Many experimental paradigms and clinical applications will require local rather than systemic injection. To be useful in these contexts, viral expression must retain a high level of specificity for PV cINs. Stereotactically guided injections typically lead to a higher number of viral particles per cell compared to systemic delivery, which may result in off-target expression. To test whether increasing the viral load altered the specificity, we locally injected the same volume of rAAV-E2-dTomato at various titers in the cortex of adult mice and assessed reporter expression within PV cINs after a week (figure 2c). The results show that while higher titers had increased levels of reporter expression, no significant alteration of specificity was observed. Targeting PV cINs at early postnatal stages has been hampered by the relatively late expression of parvalbumin (around 15 days after birth – P15) and the lack of other early markers for this population. Their involvement in developmental disorders highlights the need to target and manipulate this population during cortical circuit assembly. While complex genetic strategies offer a partial solution to achieving this in mice (i.e. Lhx6-Cre, Sst-Flp and Cre and Flp-dependent reporter), even this approach does not offer the means to easily manipulate these neurons before the second postnatal week. To test whether the E2 enhancer can target fast-spiking cINs before the onset of expression of parvalbumin, we examined its activity at various postnatal stages. To this end, we tiled our analysis across the early postnatal period, through a series of stereotactically-guided injections of rAAV-E2-dTomato (figure 2d). First, we assessed the selectivity of the reporter upon the onset of parvalbumin expression at P15. This revealed that we could obtain greater than 50% selectivity for PV cINs upon injection at P1, increasing to 67% by P7 injection and over 80% after P10 injection. We next tested if we could use this approach to label PV cINs prior to P15. To identify fast-spiking cINs in this context, we relied on Lhx6-Cre/Intact transgenic mice, in which GFP is expressed in MGE-derived interneurons (both PV cINs and SST cINs). By co-staining for SST, the PV cINs can be distinguished as GFP-positive/SST-negative. We obtained 72% and 78% specificity for PV with a P4-P7 or a P7-P10 time course, respectively. Thus, our approach provides a means to study these neurons during circuit maturation using a single viral injection.

### Viral monitoring and manipulation of PV cINs in mice

Having demonstrated the fidelity of E2-directed expression for PV cINs with differing modes of injection and across developmental stages, we then explored the utility of this vector for studying connectivity (using a presynaptic reporter) and activity (using calcium-imaging). When E2 was used to drive a synaptophysin-tdTomato fusion gene (36), reporter expression was restricted pre-synaptically to PV cINs, with terminals peri-somatically located onto pyramidal neurons (figure 3a). When this vector was used to drive GCaMP6f expression (37), we demonstrated that PV cINs were recruited upon whisker stimulation (figure 3b and supplementary figure 5a). Together these results demonstrate that E2 provides an effective means to monitor various aspects of PV cIN biology. We next examined whether E2 was sufficient to elicit functional changes in activity using chemo-or optogenetic approaches. E2 was used in adult animals to direct the expression of the chemogenetic receptor PSAM4-5HT3-LC (38). We observed that PV cINs in brain sections collected from these animals, when exposed to the actuator varenicline, could be induced to fire when current clamped below threshold (figure 3c). Similar results were obtained using the chemogenetic receptor Gq-DREADD (39 – supplementary figure 5c). Finally, both constant and high frequency laser stimulation of PV cINs expressing the red-shifted opsin C1V1 in brain slices (40) resulted in firing time-locked to the stimulus. Demonstrating that engagement of these neurons resulted in concomitant local inhibition, pyramidal neuron activity in the vicinity of virally labeled PV cINs was consistently interrupted by laser stimulation. Notably, this effect was abolished by treatment with picrotoxin (figure 3d and supplementary figure 5b). Having confirmed the efficacy of the method *ex vivo,* we next sought to explore our ability to alter excitatory networks in vivo by opto-genetically stimulating PV cINs. Three weeks following local injection of AAV-E2-C1V1 in the primary visual cortex of adult animals, single unit recordings within the infected region were performed both at baseline and upon laser stimulation. The identity of recorded neurons was distinguished based upon their spike width and maximal firing frequency. Inhibitory interneuron firing rates were increased by laser stimulation, while excitatory neuronal firing was silenced (figure 3e). Together, these data show that E2 is able to functionally engage PV cINs and elicit network inhibition using chemo or optogenetics approaches both *ex vivo* and *in vivo.*

**Figure 3.**
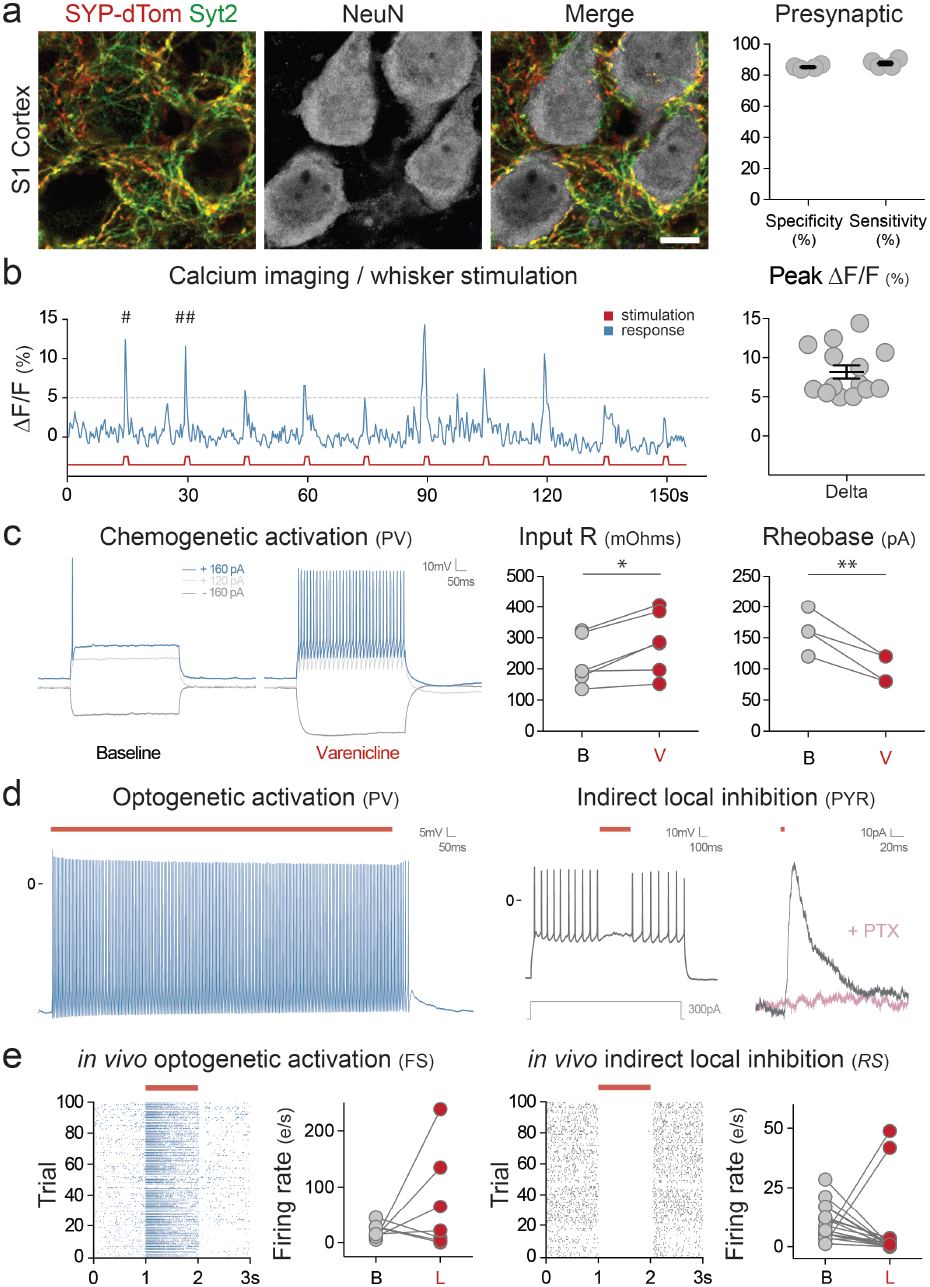
Viral monitoring and manipulation of PV cINs in mice. Mice were injected locally in the S1 cortex (**a** – P10 injection with rAAV-E2-SYP-dTomato; **b** – P14 injection with rAAV-E2-GCaMP6f; **d, e** – Adult injection with rAAV-E2-C1V1-eYFP) or systemically (**c** – Adult injection with rAAV-E2-PSAM4-5HT3-LC-GFP). (**a**) Representative images of the co-localization between the SYP-dTomato reporter and the synaptic marker Syt2 one-week post-injection and corresponding quantification. (**b**) Ca2+ imaging upon whisker stimulation was performed 2-3 weeks post-injection. On the right panel, the success rate was calculated as the proportions of ΔF/F peaks above threshold in response to whisker stimulation. (**c**) Current clamp recording was performed on brain sections 4 weeks after injection. The traces show a representative cellular response at the indicated currents at both baseline and after bath application of varenicline. (**d**) Current clamp recording was performed on brain sections 1 week after injection. Cells expressing the viral reporter were exposed to 2 seconds of constant laser stimulation (550 nm) while the voltage was recorded over 3 seconds. Neighboring pyramidal cells that did not express the viral reporter were also recorded from during laser stimulation. (**e**) In-vivo single-unit analysis of neuronal activity. Raster plots of virally infected neurons upon laser stimulation and corresponding population quantification data. The left panels show fast-spiking cells and the right panels show regular spiking excitatory cells. Notably, due to the mosaic nature of local viral injection, individual cell responses were bimodal. This presumably reflects whether or not particular cells were infected. Scale bars represent 5um. The red bars represent laser stimulation. On the graphs, dots represent individual measurements and the lines represent average +/− s.e.m. Values for specificity and sensitivity are listed in the supplementary table 2.

### Viral targeting and manipulation of PV cortical interneurons in primates including humans

The high degree of sequence conservation of the E2 enhancer across mammalian species is suggestive of a conserved role in gene regulation. We thus sought to establish whether this element could be used to target PV cINs across mammalian species. Using systemic (marmoset) or focal injections (rat and macaque) of the E2 virus, we were able to target PV cINs with approximately 90% specificity (figure 4A). It has been discovered that human brain tissue obtained during surgical resection can be cultured for prolonged periods (41). Taking advantage of this exceptional ability of human brain tissue to remain healthy *ex vivo*, we exposed freshly resected subiculum or medial temporal cortex to E2 virus. Over the two-week culture period, we observed the progressive appearance of fluorescently labeled cells. In regions where PV staining was reflective of the expected distribution of these cells, virally labeled cells were PV-positive (figure 4bi; see methods for details). In addition, the majority of cells within both the cortex and subiculum showed the characteristic hallmarks of PV INs as indicated by multiple criteria. These include morphology as well as maximum firing rate when evoked through direct depolarization or optogenetic light stimulation (figure 4Bbii-iv and supplementary figure 6). Importantly, the human version of the E2 enhancer showed the same degree of specificity for PV cINs upon injection in mice, further demonstrating that non-coding regions of the genome characterized by a high degree of sequence conservation are likely to retain their functional properties across species. Finally, truncation of both the 5’ and 3’ ends of this enhancer resulted in a drastic reduction of specificity, suggesting that the functional boundaries of the E2 enhancer have been optimally identified (supplementary figure 7). Altogether these results indicate that the E2 vector provides an effective tool for targeting and manipulating PV cINs across mammals, including humans.

**Figure 4.**
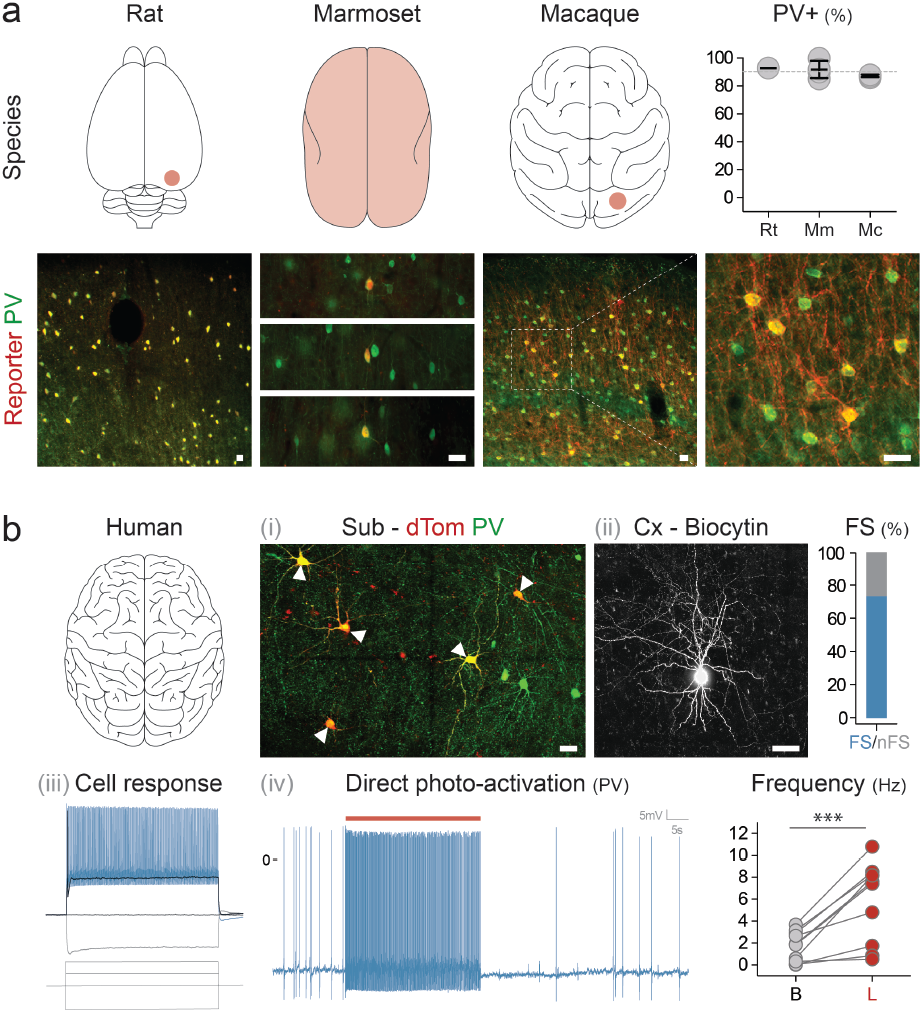
Viral targeting and manipulation of PV cINs in primates including humans. (**a**) Animals from indicated species were locally (rat and macaque) or systemically (marmoset) injected with rAAV-E2-C1V1-eYFP (macaque) or rAAV-E2-dTomato (rat and marmoset) and analyzed 2-8 weeks post-injection. (**b**) Human brain tissue obtained from surgical resection was exposed to either rAAV-E2-dTomato (**i-iii**) or rAAV-E2-C1V1-eYFP (**iv**) and maintained in culture for 7-14 days. The upper right panel shows the proportion of fast-spiking neurons among the virally labeled cells assessed by electrophysiological recordings of intrinsic properties. (**iv**) Electrophysiology current clamp recording of virally labeled cells upon laser stimulation. Scale bars represent 25um. The red bars represent laser stimulation and the arrowheads point at neurons co-expressing PV and the viral reporter. On the graphs, dots represent individual measurements and the lines represent average +/− s.e.m. Values for specificity are listed in the supplementary table 2.

### Identification of viral enhancers with regional specificity

To demonstrate that our enhancer selection method is generalizable, we identified a further 25 candidates in the vicinity of seven genes whose expression was enriched in PV cINs across species (see methods). Systemic injection of AAVs containing these sequences revealed that four of them displayed an above 90% selectivity for PV cINs. Notably, among these enhancers, the relatively few virally labeled neurons that did not express PV were positive for the pan-interneuron marker Gad1. Each of these four enhancers were specific for distinct but overlapping subsets of the PV-expressing neurons. Specifically, while E11 and E14 showed a bias for PV cINs in the upper layers of the cortex, the E22 enhancer restricted expression almost exclusively to the cortex, with only a few cells showing low levels of expression elsewhere. By contrast, the E29 virus showed the most global expression, targeting the entire population of PV-expressing neurons throughout the central nervous system. All of our enhancers have a high degree of sequence conservation and were selected from genes whose expression profile is similar across species (figure 5a). To directly test that this results in similar functionality across species, we performed local injection of the AAV-E22-dTomato in V1 of a macaque. This showed that, similarly to mouse, the expression of the viral reporter was restricted to PV cINs (figure 5b). The combination of regional selectivity and conservation of expression across species opens the possibility of using these tools for targeted therapies to correct abnormal brain function.

**Figure 5.**
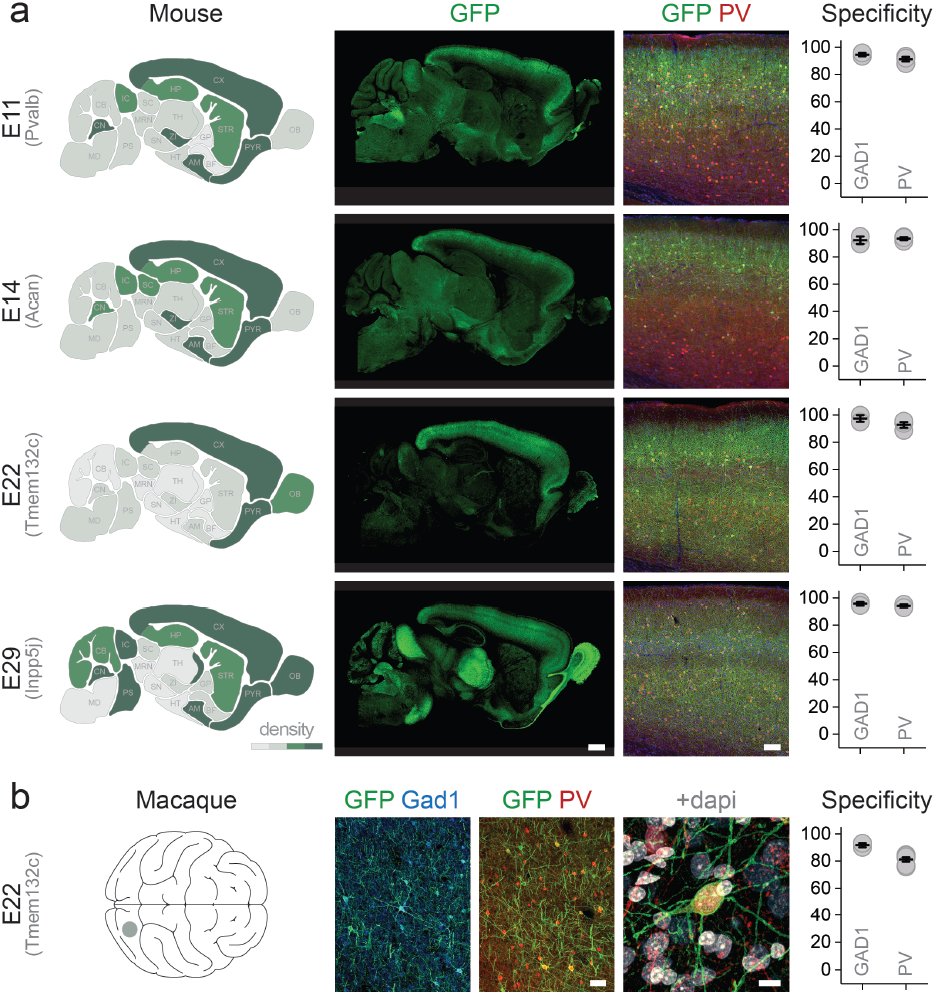
Identification of viral enhancers with regional specificity. (**a**) Adult mice were injected systemically with the indicated AAV and analyzed 3 weeks postinjection. IHC for the reporter and indicated markers in the S1 was used to assess the density of neuronal cell-bodies expressing the viral reporter (left panels) and the specificity of expression of the viral reporter for the indicated markers. (**b**) An adult macaque was injected in V1 with rAAV-E22-eGFP and analyzed 8 weeks postinjection with immunohistochemistry for the reporter and indicated markers. Scale bars represent 100 um (a), 50 um (b left) and 10 um (b right). On the graphs, dots represent individual measurements and the lines represent average +/− s.e.m. Values for specificity are listed in the supplementary table 2.

## DISCUSSION

A challenge in understanding neurological disorders stems from the complexity of the neuronal types involved. Here, in an effort to deconvolve the cellular actions of a particular disease gene, we systematically dissected the *Scn1a* genomic locus. To this end, we identified ten different enhancers distributed across the intronic and intergenic region of this gene. By creating AAVs whose expression depends on each of these enhancers, we found three that recapitulate the global pattern of *Scn1a* gene expression. As loss of expression of *Scn1a* is especially associated with PV cIN disfunction (15–19), we focused on the E2 enhancer, which was effective for selectively targeting these neurons, not only in rodents but also within various primates, including humans. This enhancer also proved useful for investigating various aspects of PV cIN function including connectivity, monitoring excitability, and for manipulating their activity with chemogenetic and optogenetic effectors. The demonstration of the utility of the E2 enhancer in a range of species, highlights its usefulness for basic and clinical applications. Notably, this approach provides the means to evaluate and compare the circuit contributions and function of PV cINs across species, regardless of their genetic accessibility (42,43).

A major impediment in examining early dynamics of circuit maturation is the inaccessibility to specific cell types without the use of transgenic animals. Young PV cINs have been particularly problematic to target even with complex genetic strategies. Given their abundance (they represent 40% of all inhibitory cINs), as well as their involvement in both the maturation of cortical circuits and neurodevelopmental disorders, gaining access to them prior to the onset of PV expression has been long awaited by the field. The specificity of the E2 enhancer at these developmental stages and the simplicity of the viral injection over the use of transgenic animals provides the means to study both their normal development and role in disease.

We began this study by examining enhancers at a specific disease locus, the *Scn1a* gene. By identifying key regulatory elements for each of the cell types that express this gene, we clarified its regulatory landscape. Many SNPs associated with the *SCN1A* locus map to intron 1 (44–46). The three enhancers we characterized were located within this region, perhaps indicating that these SNPs represent mutations affecting the expression of *Scn1a*. Corroborating this hypothesis, GTEx data show multiple eQTLs within these enhancers that are associated with alterations in *SCN1A* expression in humans (GTEx Scn1a, 47). As *Scn1a* expression is largely restricted to the PV cINs (17,18), it is tempting to speculate that mutations of the E2 enhancer may be a direct cause of Dravet syndrome.

More broadly, the enhancers identified in this study provide access to neuronal populations with particular clinical relevance. Most obviously these enhancers can potentially be leveraged to alleviate debilitating aspects of Dravet syndrome, through either gene therapy or modulation of neuronal activity (48). We demonstrated that local and systemic injections can be used for effective viral delivery to the brain. With local injections, one could possibly ameliorate focal epilepsy, prefrontal cortex dysfunction or hippocampal memory disorders. By contrast, systemic introduction of virus could be used in contexts where global interventions are necessary, for instance to correct generalized seizures, or for psychiatric and neurodegenerative disorders. Our study shows that the rigorous identification of regulatory elements provides a road map for accessing specific cell types for therapeutic contexts.

Importantly, we demonstrated that our enhancer selection method is generalizable to other genes. Overall, we found a set of 7 enhancers with unique specificity for both distinct neuronal populations and regions of the central nervous system. Even with the stringent criteria that we applied (>90% selectivity for the target population), our method has a remarkable (>20%) success rate. Moreover, as predicted by the high degree of sequence conservation, the subset of enhancers that we tested proved equally selective and effective across species including humans. Taken together our findings support that these methods provide a reliable means to systematically identify cell-type specific enhancers that work across species.

## ONLINE METHODS

### Animals

Mice. Female C57BL/6J mice (Mus musculus; 10 weeks old) were obtained from Jackson Labs (Bar Harbor, ME – stock# 000664). Rat. Sprague Dawley rats (adult 150--250 gm) were obtained from Charles River labs, Kingston, NY. Marmosets. One female common marmoset (Callithrix jacchus, 6.0 years old) was obtained from the colony at Massachusetts Institute of Technology. Macaques. One male macaque (Macaca mulatta; 15.0 years old) was obtained from the California National Primate Research Center at the University of California, Davis. All the animals were maintained in a 12 light/12 dark cycle with a maximum of five animals per cage for mice and one animal per cage. Marmosets and macaques were socially housed. All animal maintenance and experimental procedures were performed according to the guidelines established by the Institutional Animal Care and Use Committee at the Broad Institute of MIT and Harvard (mice), McGovern research institute at MIT (rats and marmosets) and Salk Institute for Biological studies (macaques).

### scATAC-seq library preparation and sequencing

Male hemizygous Dlx6a-Cre mice (Jax stock #008199) were crossed with female homozygous INTACT mice (flox-Sun1-eGFP, Jax stock #021039) to yield Dlx6a-Cre::INTACT offspring for scATAC-seq experiments. Brains from P28 Dlx6aCre::INTACT mice were harvested, sectioned coronally on a mouse brain slicer (Zivic Instruments), and regions of interest were dissected in ice-cold ACSF. Tissue was then transferred to a dounce homogenizer containing Lysis Buffer (10 mM Tris-HCl, 10 mM NaCl, 3 mM MgCl2, 0.01% Tween-20, and 0.01% IGEPAL CA-630, 0.001% Digitonin). Tissue was homogenized with 10 strokes of pestle A, 10 strokes of pestle B, and incubated for 5 min on ice before being filtered through a 30 μm filter and centrifuged at 500xg for 10 min at 4°C. The pellet was resuspended in 1% BSA for sorting for GFP+ nuclei on a Sony SH800S cell sorter. Nuclei were sorted into Diluted Nuclei Buffer (10X Genomics). scATAC-seq libraries were prepared using the Chromium Single Cell ATAC Solution (10X Genomics). Libraries were sequenced using a Nova-Seq S2 100 cycle kit (Illumina).

### scATAC analysis

Raw sequencing data were passed through the Cell Ranger ATAC pipeline (10X Genomics). The fragments files were then used to generate snap files for analysis using the snapATAC package (https://doi.org/10.1101/615179). Cells were clustered using graph-based clustering (k=15, 24 principle components). Gene activity scores were generated as described in the snapATAC package and used to determine clusters corresponding to interneuron cardinal classes. For each cardinal class, bigwig files were generated and peaks were called using macs2 for input into the Integrated Genome Browser and enhancer selection. Peaks across cardinal classes were compared using bedtools Jaccard.

### Enhancer selection

*Selection*. Candidate regulatory elements were manually curated from a list of elements generated by intersecting the “context” region (Scn1a intergenic region + intron1) with both the scATACseq peak and the conservation data from PhasCons Analysis (see below). *Accessibility.* scATAC-seq peaks obtained for each of the 4 classes of interneurons (see above) were examined. Bulk ATAC-seq data for cortical excitatory neurons generated in Mo et al. 2015 were downloaded from the GEO repository and discretized as peaks using MACS2 (https://github.com/taoliu/MACS). *Conservation.* The “phascons 60-way” track was downloaded from the UCSC portal (https://genome.ucsc.edu) in BED file format and filtered using a custom R script to remove any element smaller than 10bp and fuse any element separated by less than 50bp using Bedtools/Intersect.

### rAAV cloning and production

The enhancer sequences were amplified by PCR from mouse genomic DNA using the following primers: E1: caaagtggacagaggggagg and gtgctgttgggagtggtgga (1280 bp); E2: aatctaacatggctgctata and caattgctcagagttatttt (618 bp); E3: ataaaattttattttcctaa and gaggaaatcagctacggggc (832 bp); E4: tctgacagagcaagtcttga and tatcaaaattgtatattcag (261 bp); E5: aatgttttgatatttaggag and ttgactcttaaaatttaata (663 bp); E6: ttgtcactttgttactctac and ttaaatcttaaaattttcct (606 bp); E7: gatactgtataattaattag and cttccttctggttccttttt (2430 bp); E8: attgatctccaactttttaa and gttcatccaagtaataagag (1644 bp); E9: atctcaagtgtatgtaacat and gtctttttgttttttttttt (521 bp); E10: tattgcaaaaggaaggaatg and tcatggaaaaagaaaaaatc (547 bp). The enhancers, reporters and effectors were cloned using the Gibson Cloning Assembly Kit (NEB-E5510S) following standard procedures. Specifically, for AAV-E1:10-dTomato, we amplified the dTomato coding sequence from the plasmid Addgene # 83897; for AAV-E2-SYP-dTomato, we amplified the Synaptophysin-tdTomato coding sequence from the plasmid Addgene # 34881; for AAV-E2-GCaMP6f, we amplified the GCaMP6f coding sequence from the plasmid Addgene # 83899; for AAV-E2-C1V1-eYFP, we amplified the C1V1-eYFP coding sequence from the plasmid Addgene # 35499. The rAAVs were produced using standard production methods. PEI was used for transfection and OptiPrep gradient (Sigma, USA) was used for viral particle purification. Serotype 1 was used to produce the AAVs for local injections in mice and rats. Serotype 9 was used for systemic injection in marmosets and serotype PHPeB was used for both local injection in macaques and systemic injections in mice. Titer was estimated by qPCR with primers for the WPRE sequence that is common to all constructs. All batches produced were in the range of 10E+10 to 10E+12 viral genomes per milliliter.

### Local and systemic viral injections

Mouse local S1. Local injection in adult mice were performed by stereotactically guided injections in the somatosensory cortex with the following coordinates 1.0 mm posterior, 2.9 mm lateral, 0.7/0.45 mm ventral relative to Bregma with 150 nL of virus. Mouse systemic. For systemic injection in adult mice, approximately 10E+11 viral particles were injected in the retro-orbital sinus per animal. Postoperative monitoring was performed for five days post injection. Rat local in V1. Local injection in adult rats were performed by stereotactically guided injections in the primary visual cortex with the following coordinates 5.4 mm posterior, 4.2 mm lateral, 2.0 mm ventral relative to bregma with 670nL of virus. Marmoset systemic injection. For systemic injection in adult marmosets, approximately 10E+12 viral particles in ~ 0.7 ml of sterile PBS were injected into the saphenous vein, followed by another infusion with ~ 0.5 ml of saline. After the final infusion, pressure was applied to the injection site to ensure hemostasis. The animal was returned to its home cage and monitored closely for normal behavior post anesthesia. The animal was euthanized 51 days after viral injection. Macaque local in V1. Local injection in an adult macaque was performed by a stereotactically guided injection in the left primary visual cortex with the following coordinates: 13 mm posterior, 19 mm lateral, 23 mm superior relative to the center of the inter-aural line (based on the animal’s MRI). A total of volume of 1332 nL was injected, equally divided in 4 depths (i.e. 1.8, 1.3, 0.8 and 0.3 mm from the cortical surface).

### Electrophysiological recordings in mice

*Slice preparation for 2 to 6-weeks-old mice.* Virally injected mice were anesthetized with isofluorane. Upon loss of reflexes, mice were transcardially perfused with ice-cold oxygenated ACSF containing the following (in mM): 87 NaCl, 75 sucrose, 2.5 KCl, 1.25 NaH2PO4, 26 NaHCO3, 10 glucose, 1 CaCl2 and 2 MgCl2. Mice were then decapitated and 300-μm thick coronal slices were sectioned using a Leica VT-1200-S vibratome and incubated in a holding chamber at 32–35 °C for 15–30 min followed by continued incubation at room temperature 20–23.5 °C (68–74 °F) for at least 45-60 min before physiological recordings. Slice containing the injection site were transferred in a recording chamber submerged with oxygenated ACSF containing the following (in mM): 125 NaCl, 2.5 KCl, 1.25 NaH2PO4, 26 NaHCO3, 10 glucose, 2 CaCl2 and 1 MgCl2 (pH = 7.4, bubbled with 95% O2 and 5% CO2). *Slice preparation for 6-weeks-old and older mice.* Acute coronal brain slices were prepared as follows. Mice were anesthetized with Avertin solution (20 mg/ml, 0.5 mg/g body weight) and transcardially perfused with 15 to 20 ml of ice-cold carbogenated (95% O2, 5% CO2) cutting solution containing the following: 194 mM sucrose, 30 mM NaCl, 4.5 mM KCl, 1.2 mM NaH2PO4, 0.2 mM CaCl2, 2 mM MgCl2, 26 mM NaHCO3, and 10 mM D-(+)-glucose (with osmolarity of 340–350 mOsm). The brains were then rapidly removed and placed in ice-cold cutting solution for slice preparation. Coronal slices (300 μm) were prepared and then incubated at 32°C with carbogenated artificial cerebral spinal fluid (aCSF) for 10 to 15 minutes. The slices were then incubated at room temperature for at least 1 hour in aCSF that contained the following: 119 mM NaCl, 2.3 mM KCl, 1.0 mM NaH2PO4, 26 mM NaHCO3, 11 mM glucose, 1.3 mM MgSO4, and 2.5 mM CaCl2 (pH 7.4, with osmolarity of 295–305 mOsm) at room temperature for at least 1 hour. Current clamp. For interneuron recording, 10 μM CNQX, 25 μM AP-5 and 10 μM SR-95531 were also added to block AMPA, NMDA and GABA-A receptors, respectively, to measure the cell-intrinsic effect of optogenetic and chemogenetic stimulation. Whole-cell current-clamp recordings were obtained from visually identified cells expressing the viral reporter using borosilicate pipettes (3–5 MΩ) containing (in mM): 130 K-gluconate, 6.3 KCl, 0.5 EGTA, 10 HEPES, 4 Mg-ATP, 0.3 Na-GTP and 0.3% biocytin (pH adjusted to 7.3 with KOH). Upon break-in, series resistance (typically 15–25 MΩ) was compensated and only stable recordings (<20% change) were included. Data were acquired using a MultiClamp 700B amplifier (Molecular Devices), sampled at 20 kHz and filtered at 10 kHz. All cells were held at −60 mV with a DC current, and current-step protocols were applied to obtain firing patterns and to extract basic sub-threshold and supra-threshold electro-physiological properties. Voltage clamp. Cells not expressing the viral reporter were selected according to their pyramidal-cell-shaped soma under IR-DIC visualization and recorded with pipettes containing (in mM): 130 Cs-gluconate, 0.5 EGTA, 7 KCl, 10 HEPES, 4 Mg-ATP, 0.3 Na-GTP, 5 phosphocreatine, 5 QX-314 and 0.3% biocytin (pH adjusted to 7.3 with CsOH). Cells were held continuously at 0 mV for baseline and optogenetic or chemogenetic stimulation. For both current and voltage clamp recording, a baseline of at least 2 min was recorded before stimulation. Small pulses (−20 pA or −5 mV, 100 ms at 0.2 Hz or 0.5 Hz) were applied throughout the baseline and CNO application to monitor series resistance changes. Data were analyzed offline using Clampfit 10.2 software (Molecular Devises).

### *In-vivo* calcium imaging

Approximately 100 nL of AAV-E2-GCaMP6 virus was injected in barrel cortex at postnatal day 10. At P27-P34, craniotomies were implanted over the injection site and widefield calcium imaging performed after recovery from craniotomy procedure. Briefly, anesthetized (1.5% isoflurane) mice were imaged at 3-4Hz with 4x magnification (Thorlabs CCD camera – 1501M-USB, Thorlabs LED stimulation – DC4104), while air puffs (100-200ms duration, Picospritzer III) at specific intervals (5-20s) were directed at contralateral whiskers. Multiple recordings were performed and afterward the mouse was perfused for histological analysis. Recordings were analyzed in ImageJ by calculating the F/F (change in fluorescence/average fluorescence) for each recording and synched whisker stimulation. A threshold of (5%) F/F was set for both stimulated and spontaneous calcium signal response.

### Electrophysiological recordings in human

*Tissue preparation, culture protocol and inoculation of virus.* Four participants (2 males / 2 females; age range 22-57 years) underwent a surgical procedure in which brain tissue (temporal lobe and hippocampus) was resected for the treatment of drug resistant epilepsy. In all cases, each participant had previously undergone an initial surgery for placement of subdural and/or depth electrodes for intracranial monitoring in order to identify the location of seizure onset. The NINDS Institutional Review Board (IRB) approved the research protocol (ClinicalTrials.gov Identifier NCT01273129), and we obtained informed consent from the participants for experimental use of the resected tissue. 300 um slices from both hippocampus and temporal lobe were attained (Leica 1200S Vibratome; Leica Microsystems, Bannockburn, IL) in ice-cold oxygenated sucrose based cutting solution (100 mM sucrose, 80 mM NaCl, 3.5 mM KCl, 24 mM NaHCO3, 1.25 mM NaH2PO4, 4.5 mM MgCl2, 0.5 mM CaCl2, and 10 mM glucose, saturated with 95% O2 and 5% CO2) within 30 minutes following neurosurgical resection. Slices were then incubated in the sucrose cutting solution at 33°C for 30 minutes and allowed to cool to room temperature for 15-30 minutes. The slices were transferred to culture media (Eugène et al., 2014) and placed in an incubator (5% CO2) at 35oC for 15 minutes of equilibration. Each individual slice was then transferred onto a 30 mm Millicell Cell Culture Insert (Millipore; Cat No. PICM0RG50) for interface culture and incubated as above. After 12 hours, the culture medium was changed and 1-2 ul of pAAV-S5E2-dTomato with or without pAAV-S5E2-C1V1-eYFP was directly pipetted onto each slice and placed back into the incubator. For hippocampal slices, the virus was targeted to the subiculum subfield. Culture medium was routinely changed every 2-3 days until electro-physiological analyses. Electro-physiological recordings. Electrophysiological recordings from cultured human slices were performed between 7 to 14 days after viral inoculation. Cultured human slices were transferred to a recording chamber perfused with extracellular solution (130 mM NaCl, 3.5 mM KCl, 24 mM NaHCO3, 1.25 mM NaH2PO4-H2O, 10 mM glucose, 2.5 mM CaCl2 and 1.5 mM MgCl2 saturated with 95% O2/5% CO2 (pH 7.4; 300-310 mOsm) at a rate of 3 – 4 ml/min at 33°C. Whole cell patch clamp recordings from pAAV_S5E2-dTomato or pAAV-S5E2-C1V1-eYFP infected neurons were performed with an intracellular solution of the following composition: 130 mM K-gluconate, 10 mM HEPES, 0.6 mM EGTA, 2 mM MgCl2, 2 mM Na2ATP, 0.3 mM NaGTP and 0.5% biocytin (pH adjusted to 7.4; osmolarity adjusted to 285 – 300 mOsm). In some recordings, 130mM K-gluconate was replaced by 90 mM K-gluconate/40 KCl. Intrinsic membrane and firing properties were assayed essentially as described previously. 550nm light stimulated optogenetic activation of C1V1 was delivered to the slices via the 40X water immersion objective using a CoolLED pE-4000 Illumination system (Andover, UK). Biocytin reconstruction and immunocytochemistry. After electrophysiological recording slices were drop-fixed in 4% paraformaldehyde in 0.1M PB overnight. Slices were washed in 0.1M PB (3 × 15 minutes) and permeabilized / blocked in 0.5% Triton X-100/10% goat serum in 0.1M PB for at least 2 hours at room temperature. For combined biocytin recovery and immunocytochemistry an initial incubation (4°C for 40 hours) in primary antibodies diluted at 1:1000 was performed (rabbit anti-PV, Abcam Cat No: ab11427; guinea-pig anti-RFP, SYSY, Cat No: 390005). Slices were washed in 0.1M PB at room temperature 4 × 30 minutes and incubated in secondary antibodies (1:1000 for goat anti guinea-pig Alex-flour 555, Thermofisher Cat No. A21435; 1:500 for goat anti-rabbit Alexa-flour 647, Thermofisher Cat No. A32733 and 1:1000 Streptavidin Alexa FluorTM 488; Thermofisher S1123) overnight at 4oC. After a final wash procedure (4 × 30 minutes) the slices were mounted on microscope slides with Prolong Gold antifade (Thermofisher; Cat No. P36930) for subsequent confocal microscopy.

### Immunohistochemistry

Citations with validation data for each antibody are reported on the providers' websites. All animals injected with the virus were transcardially perfused with 4% paraformaldehyde (PFA). The brains were placed in 4% PFA overnight then sectioned at 50 μm using a Leica VTS1000 vibrosector. Floating sections were permeabilized with 0.1% Triton X-100 and PBS for 30 min, washed three times with PBS and incubated in blocking buffer (5% normal donkey serum in PBS) for 30 minutes. The sections were then incubated overnight in blocking buffer with the indicated combination of the following primary antibodies at 4 °C: chicken anti-GFP at 1:1,000 (Abcam USA, ab13970); rabbit anti-DsRed at 1:1,000 (Clontech USA 632496); goat anti-PV at 1:1,000 (Swant USA, PVG-213); guinea-pig anti-PV at 1:1,000 (Swant USA, GP-72); rabbit anti-SST at 1:2000 (Peninsula USA, T-4103.0050); mouse anti-Synaptotagmin-2 at 1:250 (ZFIN USA, #ZDB-ATB-081002-25). The sections were then washed three times with PBS, incubated with Alexa Fluor-conjugated secondary antibodies at 1:1000 (Invitrogen, USA), counterstained with DAPI (Sigma, USA) and mounted on glass slides using Fluoromount-G (Sigma, USA). The staining of PV IHC within human brain tissues was highly variable. As such, estimates of viral specificity were made within regions of cortex and subiculum where staining density was reflective of the known distribution and density of these cells. Given this variability, accurate quantitation was not possible.

### *In-situ* hybridization

The in-situ hybridization probes (Gad1; product #400951, Pvalb; product #421931, VIP; product #415961) used were designed by Advanced Cell Diagnostics (Newark, CA, USA). The reagents in the RNAscope^®^ Multiplex Fluorescent Reagent Kit v2 (product # 323100), RNAscope^®^ Probe Diluent (product #300041), HybEZ™ oven (product #321710/321720), humidity control tray (product # 310012), and HybEZ Humidifying Paper (product #310025) were also from Advanced Cell Diagnostics. TSA Plus Fluorescein, TSA Plus Cyanine 3, and TSA Plus Cyanine 5 from PerkinElmer (#NEL741, #NEL744, and #NEL745). Brain tissue was processed as mentioned in the immunohistochemistry section. Brain sections were washed one time in PBS followed by three washes in 0.1% Triton X-100 and PBS, mounted on Superfrost Plus glass slides (Fisher Scientific, 12-550-15) and baked at 60°C in the HybEZ oven for 25 minutes. The slides were then submerged in 4% PFA for 30 minutes then washed 3 times in H2O. RNAscope H2O2 was applied to each section for 5 minutes at room temperature. The slides were then washed 3 times in H2O before being submerged in pre-warmed 90°C H2O for 15 seconds and followed by pre-warmed 90C RNAscope Target Retrieval for 15 minutes. Slides were washed 3 times in H2O before RNAscope Protease III was applied onto each section and then incubated for 15 minutes at 40°C in the HybEZ oven. Slides were washed 3 times in H2O and then incubated with probe solution diluted to 1:50 with probe diluent for 2 hours at 40°C in HybEZ oven. Next, the sections were washed three times in RNAscope wash buffer followed by fluorescence amplification. Of note, probes against the RNA of the reporter revealed a non-specific staining that we speculate comes from the viral DNA. In order to reveal the viral reporter, we followed the RNAscope protocol with an IHC amplification of the dTomato. The sections were incubated in blocking solution (0.3% Triton X-100 plus 5% normal horse serum in PBS) for 30 minutes. Then sections were incubated in antibody solution (0.1% Triton X-100 plus 5% normal horse serum in PBS) with rabbit anti-DsRed at 1:250 (Clontech USA 632496) at 4°C overnight. The sections were then washed three times with PBS, incubated with Alexa Fluor-conjugated secondary antibodies at 1:500 (Invitrogen, USA), counterstained with DAPI (Sigma, USA) and mounted on glass slides using Fluoromount-G (Sigma, USA).

### Quantifications and statistics

For strength of expression, fluorescence images were taken at a standardized magnification and exposure time and the average pixel intensity of the cells bodies of each cell expressing the viral reporter was recorded and reported as an average over all cells per enhancer. For quantification of colocalization, cells expressing the indicated reporter were counted using only the corresponding color channel, and then among these cells, the number of cells coexpressing the marker of interest were counted. A cell was considered to be positive for a given marker if the corresponding signal was above background fluorescence. The ratio of cells coexpressing both markers over the total number of cells expressing only the reporter was then calculated, reported here as mean ± s.e.m. Quantifications were performed using a minimum of two independent biological replicates (see supplementary table 2). Several sections from the same animal were used when indicated. Data collection and analysis were not performed blind to the conditions of the experiments, but experimenters from different research groups performed the quantification. No statistical methods were used to predetermine sample sizes, but our sample sizes are similar to those reported in previous publications.

## DATA AVAILABILITY

The data that support the findings of this study are available from the corresponding author upon reasonable request. The scATAC-seq dataset will be available on GEO repository and the AAV plasmids will be available on Addgene upon publication of this work in a peer-reviewed journal.

## ACKNOWLEDGEMENT

J.D is supported by NIH grants R01-MH111529 and UG3MH120096, as well as support from the Simons Foundation Award 566615 and a gift from the Friends-Of-FACES foundation. T.P.F. is supported by fellowships from the Belgian American Educational Foundation and the George E. Hewitt Foundation for Medical Research, a NARSAD Young Investigator Grant from the Brain & Behavior Research Foundation and a Kavli Institute for Brain and Mind Innovative Research Grant. IRP of NINDS and Eunice Kennedy Shriver NICHD Intramural Grant were awarded to CJM. JS is supported by National Institute of Neurological Disorders and Stroke Grant No. K99 NS106528. G.F. is supported by NINDS grants NS081297, NS074972, NIMH grant MH071679 and NIH grant UG3MH120096, the Harvard’s Dean Initiative, as well as support from the Simons Foundation Award 566615. We are indebted to all of patients who have selflessly volunteered to participate in this study.

## AUTHOR CONTRIBUTION

DV and JDL designed and performed experiments, analyzed data, prepared figures and wrote the manuscript. KA performed and analyzed the scATAC-seq experiments and GS provided computational support for the analysis. KAP, RC, XY and KAZ performed experiments and analyzed data related to human brain tissue. BG, MA, SS, GS, OS, SV, LAI, KJM, ES, SH, EF, TB, IV and VS performed experiments, analyzed data related to mouse. QX and LG produced the adeno-associated in NYUAD using plasmids conceived and generated at the Broad Institute. JS and QZ performed experiments and analyzed data related to rats and marmosets. TPF and JS performed experiments and analyzed data related to macaques. OD, BLS, RB, JR, GF, ZF, CJM, GJF helped with the study design and provided collaborative support to the work. JD designed the study, conceived the viral construct, and contributed to writing the manuscript, prepared the figures and supervised the project. All authors edited and approved the manuscript.

