## Supplementary figures 1-7 and supplementary tables 1-2 for "Viral manipulation of functionally distinct neurons from mice to humans"

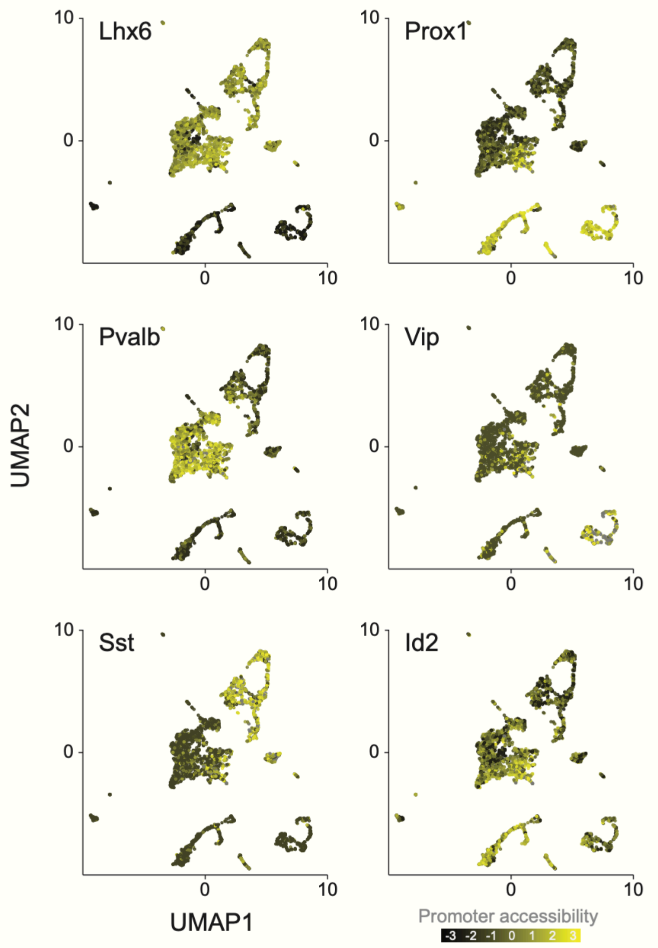


**Supplementary Figure 1.** UMAP plot of 3500 neuronal nuclei collected from 4 Dlx6a$^{Cre}$::Sun1-GFP mice colored promoter accessibility of the indicated canonical interneuron markers.

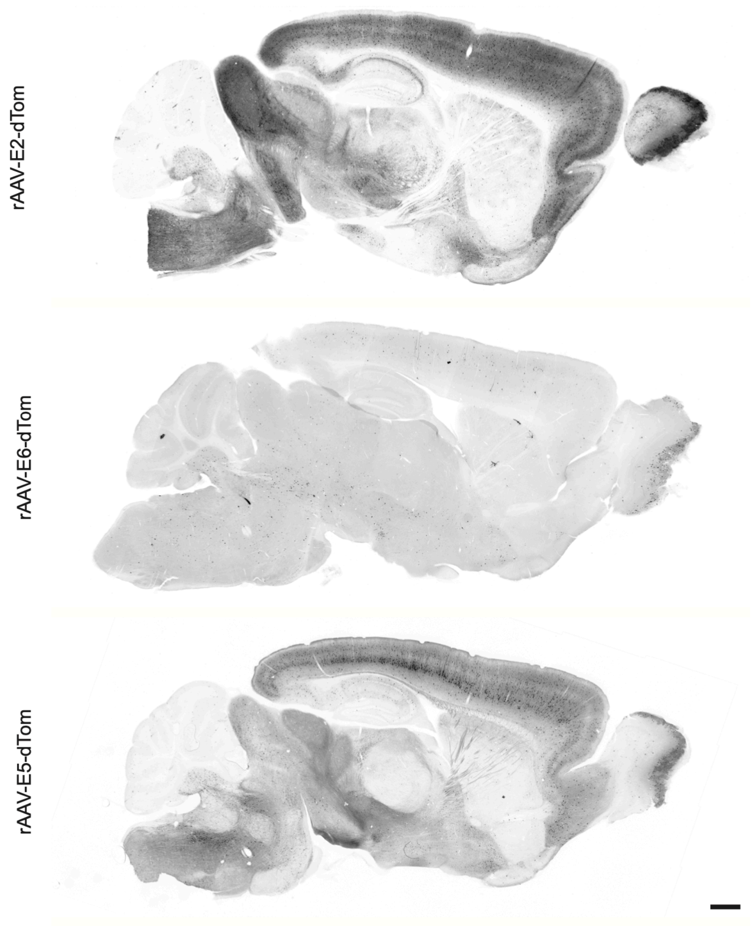


**Supplementary Figure 2.** Fluorescent images of sagittal sections from adult mice that were injected systemically with the indicated rAAV-E[x]-dTom and analyzed 3 weeks post-injection with IHC for the viral reporter. Scale bar represents 500um.

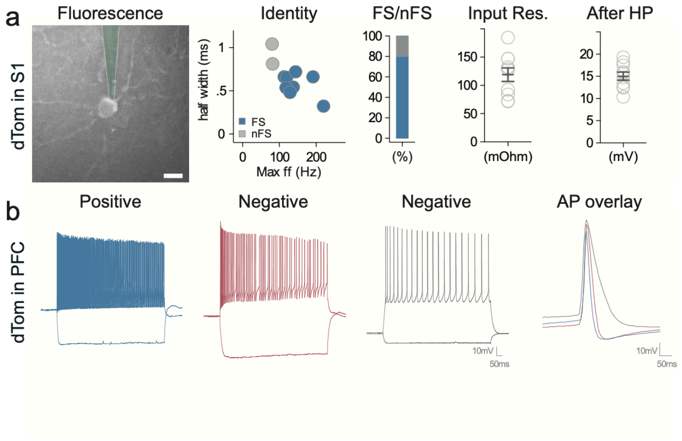


**Supplementary Figure 3.** Adult mice were injected systemically with rAAV-E2-dTomato. **(a)** Slice recording of the intrinsic properties of virally labeled neurons. The left panel shows a representative cell expressing the viral reporter. The green trace represents the recording pipet. The quantifications show the indicated parameters. The blue dots represent cells with stereotypical fast-spiking properties. **(b)** Representative slice recording traces of positive fast-spiking cell and negative. Scale bars represent 20um. On the graphs, dots represent individual measurements and the lines represent average +/- s.e.m.

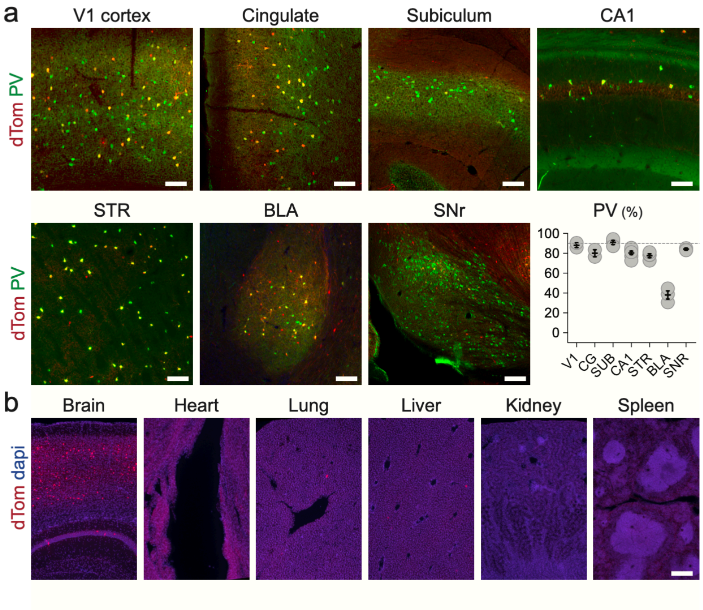


**Supplementary Figure 4.** Adult mice were injected systemically with rAAV-E2-dTomato and analyzed 3 weeks post-injection. **(a)** Coronal and sagittal sections were analyzed with IHC for the viral reporter and PV and the specificity to PV was reported across brain regions. **(b)** The native viral expression was analyzed from the indicated organs. Scale bars represent 100um (a) and 250um (b). On the graphs, dots represent individual measurements and the lines represent average +/- s.e.m. Values for specificity are listed in the supplementary table 2.

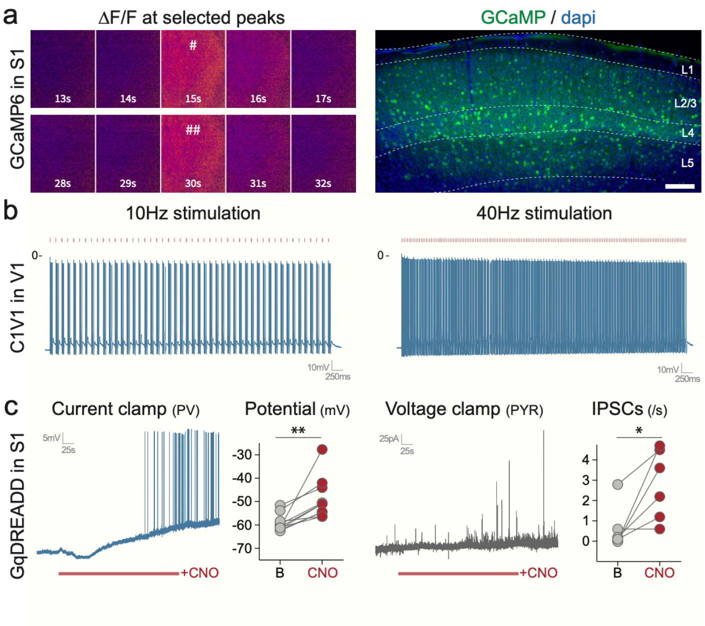


**Supplementary Figure 5.** Mice were injected systemically (**a** - P14 injection with rAAV-E2-GCaMP6f) and locally in the somatosensory cortex (**b** – rAAV-E2-C1V1-eYFP; **c** – rAAV-E2-GqDREADD). **(a)** Mice were analyzed 1-week post-injection. The left panel shows widefield images of two representative peaks shown by the pound sign in figure 3. The right panel shows a fluorescent image taken after GCaMP recordings. **(b)** Slice electrophysiology current clamp recording were performed 1-week post-injection. Cells expressing the viral reporter were targeted with either 10Hz or 40Hz laser stimulation (550nm) while the voltage was recorded over 3 seconds. **(c)** Slice electrophysiology current clamp recording were performed 1-week post-injection. The voltage was recorded before and after bath application of CNO. Scale bars represent 500um. The red bars represent laser stimulation. On the graphs, dots represent individual measurements.

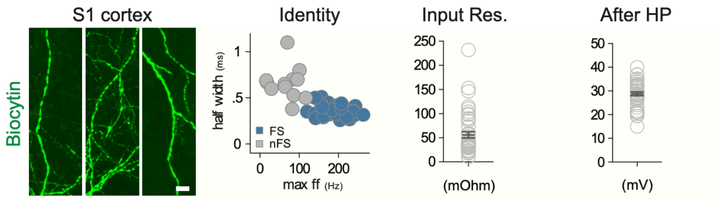


**Supplementary Figure 6.** Human brain tissue obtained from surgical resection that was exposed to rAAV-E2-dTomato and maintained in culture for 7-14 days. **(a)** Representative image of the dendrites of virally labeled cells filled with Biocytin during the recording session. **(b)** Slice recording of the intrinsic properties of virally labeled neurons. The quantifications show the indicated parameters. The blue dots represent cells with stereotypical fast-spiking properties. Scale bars represent 100um. On the graphs, dots represent individual measurements and the lines represent average +/- s.e.m.

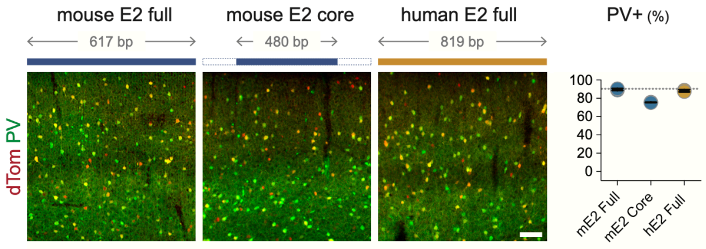


**Supplementary Figure 7.** Adult mice were injected with the indicated modified rAAV-E2-dTomato construct and analyzed 3 weeks post-injection with IHC for the viral reporter and PV. The corresponding specificity is shown in the right panel. Scale bars represent 2um. On the graphs, dots represent individual measurements and the lines represent average +/- s.e.m. Values for specificity are listed in the supplementary table 2.

| RE | Gene | Target | Specificity | Position | Chr | Start | Stop | Size (bp) | scATAC PV | scATAC SST | scATAC VIP | scATAC ID2 | bATAC Ex | Ms/Hm |
|---|---|---|---|---|---|---|---|---|---|---|---|---|---|---|
| E1 | Scn1a | PV | 22% | intergenic | chr2 | 66256056 | 66257335 | 1279 | 1 | 1 | 1 | 1 | 1 | 69% |
| E2 | Scn1a | PV | 90% | intronic | chr2 | 66364036 | 66364653 | 617 | 1 | 0 | 0 | 1 | 0 | 71% |
| E3 | Scn1a | PV | 57% | intronic | chr2 | 66383190 | 66384021 | 831 | 1 | 1 | 1 | 1 | 1 | 67% |
| E4 | Scn1a | PV | 14% | intronic | chr2 | 66387764 | 66388024 | 260 | 0 | 0 | 0 | 0 | 0 | 78% |
| E5 | Scn1a | PYR | 20% | intronic | chr2 | 66392447 | 66393109 | 662 | 1 | 1 | 1 | 1 | 1 | 72% |
| E6 | Scn1a | VIP | 88% | intronic | chr2 | 66401767 | 66402372 | 605 | 0 | 0 | 0 | 0 | 0 | 72% |
| E7 | Scn1a | PV | 33% | intronic | chr2 | 66407834 | 66410263 | 2429 | 1 | 1 | 1 | 1 | 1 | 74% |
| E8 | Scn1a | PV | 61% | intronic | chr2 | 66439814 | 66441457 | 1643 | 1 | 1 | 1 | 1 | 1 | 75% |
| E9 | Scn1a | PV | 47% | intergenic | chr2 | 66441748 | 66442268 | 520 | 1 | 0 | 0 | 0 | 1 | 72% |
| E10 | Scn1a | PV | 23% | intergenic | chr2 | 66450594 | 66451140 | 546 | 1 | 1 | 1 | 1 | 1 | 75% |
| E11 | Pvalb | PV | 90% | intronic | chr15 | 78204152 | 78204655 | 503 | 1 | 1 | 1 | 1 | 0 | 78% |
| E12 | Pvalb | PV | 59% | intronic | chr15 | 78204583 | 78204784 | 201 | 1 | 1 | 0 | 0 | 0 | 74% |
| E13 | Pvalb | PV | 67% | intronic | chr15 | 78205234 | 78205766 | 532 | 0 | 1 | 0 | 0 | 0 | 73% |
| E14 | Acan | PV | 94% | intergenic | chr7 | 79052127 | 79052622 | 495 | 1 | 0 | 0 | 0 | 0 | 72% |
| E15 | Acan | PV | 79% | intergenic | chr7 | 79053118 | 79053435 | 317 | 1 | 1 | 1 | 1 | 1 | 84% |
| E16 | Acan | PV | 58% | intronic | chr7 | 79056553 | 79057054 | 501 | 0 | 0 | 0 | 0 | 0 | 82% |
| E17 | Acan | PV | 54% | intronic | chr7 | 79079999 | 79080472 | 473 | 1 | 0 | 0 | 0 | 0 | 86% |
| E18 | Tmem132c | PV | 57% | intronic | chr5 | 127243448 | 127244121 | 673 | 1 | 1 | 0 | 0 | 0 | 70% |
| E19 | Tmem132c | PV | 57% | intronic | chr5 | 127257256 | 127257594 | 338 | 0 | 1 | 0 | 0 | 0 | 78% |
| E20 | Tmem132c | PV | 66% | intronic | chr5 | 127290515 | 127291016 | 501 | 1 | 0 | 0 | 0 | 0 | 77% |
| E21 | Tmem132c | PV | 71% | intronic | chr5 | 127300767 | 127301107 | 340 | 0 | 0 | 0 | 0 | 0 | 74% |
| E22 | Tmem132c | PV | 94% | intronic | chr5 | 127305150 | 127305592 | 442 | 1 | 0 | 0 | 0 | 0 | 75% |
| E23 | Tmem132c | PV | 64% | intronic | chr5 | 127323924 | 127324468 | 544 | 1 | 1 | 1 | 1 | 0 | 85% |
| E24 | Tmem132c | PV | 82% | intronic | chr5 | 127331966 | 127332522 | 556 | 1 | 1 | 1 | 1 | 0 | 74% |
| E25 | Tmem132c | PV | 73% | intronic | chr5 | 127355818 | 127356133 | 315 | 1 | 0 | 0 | 0 | 0 | 77% |
| E26 | Lrrc38 | PV | 72% | intergenic | chr4 | 143348892 | 143349749 | 857 | 1 | 1 | 1 | 0 | 0 | 70% |
| E27 | Lrrc38 | PV | 66% | intronic | chr4 | 143361408 | 143362362 | 954 | 1 | 0 | 1 | 0 | 0 | 71% |
| E28 | Inpp5j | PV | 83% | intergenic | chr11 | 3504821 | 3505244 | 423 | 1 | 1 | 1 | 1 | 1 | 77% |
| E29 | Inpp5j | PV | 94% | intergenic | chr11 | 3509025 | 3509652 | 627 | 1 | 1 | 0 | 0 | 0 | 74% |
| E30 | Mef2c | PV | 77% | intergenic | chr13 | 83503268 | 83504033 | 765 | 0 | 0 | 0 | 0 | 0 | - |
| E31 | Mef2c | PV | 63% | intronic | chr13 | 83507235 | 83507457 | 222 | 0 | 0 | 0 | 0 | 0 | 68% |
| E32 | Mef2c | PV | 70% | intronic | chr13 | 83515122 | 83515409 | 287 | 0 | 0 | 0 | 0 | 0 | 70% |
| E33 | Mef2c | PV | 82% | intronic | chr13 | 83518268 | 83519179 | 911 | 1 | 1 | 1 | 1 | 1 | - |
| E34 | Pthlh | PV | 48% | intronic | chr6 | 147263395 | 147263584 | 189 | 1 | 1 | 1 | 1 | 1 | 68% |
| E35 | Pthlh | PV | 86% | intergenic | chr6 | 147266874 | 147267390 | 516 | 0 | 0 | 0 | 0 | 0 | 69% |

**Supplementary Table 1.** Table containing the specifications for all tested enhancers that includes their associated gene, target population, specificity for target population, location, presence of ATAC peaks, and conservation with the human sequence.

| F | Quantification | n | mean | s.e.m |
|---|---|---|---|---|
| 1e | E1_Reporter_layer1-6 | 3 | 17.1 | 4.7 |
| 1e | E2_Reporter_layer1-6 | 3 | 78.7 | 11.4 |
| 1e | E3_Reporter_layer1-6 | 3 | 49.1 | 5.5 |
| 1e | E4_Reporter_layer1-6 | 3 | 16 | 2.6 |
| 1e | E5_Reporter_layer1-6 | 3 | 38.3 | 17 |
| 1e | E6_Reporter_layer1-6 | 3 | 8.1 | 0.2 |
| 1e | E7_Reporter_layer1-6 | 3 | 10.4 | 1.4 |
| 1e | E8_Reporter_layer1-6 | 3 | 41.9 | 9.2 |
| 1e | E9_Reporter_layer1-6 | 3 | 28.2 | 5.6 |
| 1e | E10_Reporter_layer1-6 | 3 | 22.3 | 7.2 |
| 1e | Dlx_Reporter_layer1-6 | 3 | 100.1 | 7.9 |
| 1e | E1_Reporter / Gad1_layer1-6 | 2 | 95.1 | 4.3 |
| 1e | E2_Reporter / Gad1_layer1-6 | 2 | 97.8 | 0 |
| 1e | E3_Reporter / Gad1_layer1-6 | 2 | 95 | 2.8 |
| 1e | E4_Reporter / Gad1_layer1-6 | 2 | 97.6 | 2.4 |
| 1e | E5_Reporter / Gad1_layer1-6 | 2 | 63.7 | 1.5 |
| 1e | E6_Reporter / Gad1_layer1-6 | 2 | 98.7 | 1.3 |
| 1e | E7_Reporter / Gad1_layer1-6 | 2 | 89.5 | 4 |
| 1e | E8_Reporter / Gad1_layer1-6 | 2 | 85.4 | 0.5 |
| 1e | E9_Reporter / Gad1_layer1-6 | 2 | 98.6 | 1.4 |
| 1e | E10_Reporter / Gad1_layer1-6 | 2 | 96.7 | 2.4 |
| 1e | Dlx_Reporter / Gad1_layer1-6 | 4 | 93.4 | 1.3 |
| 1e | E1_Reporter / PV_layer1-6 | 6 | 50.3 | 5.6 |
| 1e | E2_Reporter / PV_layer1-6 | 6 | 74 | 2.5 |
| 1e | E3_Reporter / PV_layer1-6 | 4 | 45.5 | 2.1 |
| 1e | E4_Reporter / PV_layer1-6 | 6 | 30.7 | 3.8 |
| 1e | E5_Reporter / PV_layer1-6 | 6 | 22.8 | 2.4 |
| 1e | E6_Reporter / PV_layer1-6 | 2 | 16 | 2 |
| 1e | E7_Reporter / PV_layer1-6 | 6 | 50.8 | 2.1 |
| 1e | E8_Reporter / PV_layer1-6 | 4 | 47.5 | 3.4 |
| 1e | E9_Reporter / PV_layer1-6 | 6 | 52.8 | 6.3 |
| 1e | E10_Reporter / PV_layer1-6 | 4 | 29 | 2.7 |
| 1f | E2_Reporter / Gad1_layer1 | 3 | 0 | 0 |
| 1f | E2_Reporter / Gad1_layer2/3 | 3 | 94.7 | 0.9 |
| 1f | E2_Reporter / Gad1_layer4 | 3 | 100 | 0 |
| 1f | E2_Reporter / Gad1_layer5 | 3 | 97.1 | 1 |
| 1f | E2_Reporter / Gad1_layer6 | 3 | 97.2 | 2.8 |
| 1f | E2_Reporter / PV_layer1 | 3 | 0 | 0 |
| 1f | E2_Reporter / PV_layer2/3 | 3 | 86.5 | 2.8 |
| 1f | E2_Reporter / PV_layer4 | 3 | 93.5 | 1.7 |
| 1f | E2_Reporter / PV_layer5 | 3 | 87.3 | 2.1 |
| 1f | E2_Reporter / PV_layer6 | 3 | 87.3 | 2.1 |
| 1f | E2_Reporter / SST_layer1 | 3 | 0 | 0 |
| 1f | E2_Reporter / SST_layer2/3 | 3 | 0 | 0 |
| 1f | E2_Reporter / SST_layer4 | 3 | 6.1 | 0.8 |
| 1f | E2_Reporter / SST_layer5 | 3 | 3.3 | 1 |
| 1f | E2_Reporter / SST_layer6 | 3 | 7.6 | 1 |
| 1f | E2_Reporter / VIP_layer1 | 3 | na | na |
| 1f | E2_Reporter / VIP_layer2/3 | 3 | na | na |
| 1f | E2_Reporter / VIP_layer4 | 3 | na | na |
| 1f | E2_Reporter / VIP_layer5 | 3 | na | na |

| F | Quantification | n | mean | s.e.m |
|---|---|---|---|---|
| 1f | E2_Reporter / VIP_layer6 | 3 | na | na |
| 1f | E5_Reporter / Gad1_layer1 | 3 | 91.7 | 8.3 |
| 1f | E5_Reporter / Gad1_layer2/3 | 3 | 98.9 | 1.1 |
| 1f | E5_Reporter / Gad1_layer4 | 3 | 95.7 | 0 |
| 1f | E5_Reporter / Gad1_layer5 | 3 | 35 | 1.7 |
| 1f | E5_Reporter / Gad1_layer6 | 3 | 94.7 | 5.3 |
| 1f | E5_Reporter / PV_layer1 | 3 | 0 | 0 |
| 1f | E5_Reporter / PV_layer2/3 | 3 | 36.3 | 7.7 |
| 1f | E5_Reporter / PV_layer4 | 3 | 15.8 | 3.8 |
| 1f | E5_Reporter / PV_layer5 | 3 | 17.8 | 4.5 |
| 1f | E5_Reporter / PV_layer6 | 3 | 12.9 | 2.1 |
| 1f | E5_Reporter / SST_layer1 | 3 | 0 | 0 |
| 1f | E5_Reporter / SST_layer2/3 | 3 | 7.1 | 4.3 |
| 1f | E5_Reporter / SST_layer4 | 3 | 22.4 | 10.3 |
| 1f | E5_Reporter / SST_layer5 | 3 | 13.1 | 4.3 |
| 1f | E5_Reporter / SST_layer6 | 3 | 12.3 | 12.3 |
| 1f | E5_Reporter / VIP_layer1 | 3 | na | na |
| 1f | E5_Reporter / VIP_layer2/3 | 3 | na | na |
| 1f | E5_Reporter / VIP_layer4 | 3 | na | na |
| 1f | E5_Reporter / VIP_layer5 | 3 | na | na |
| 1f | E5_Reporter / VIP_layer6 | 3 | na | na |
| 1f | E6_Reporter / Gad1_layer1 | 3 | 0 | 0 |
| 1f | E6_Reporter / Gad1_layer2/3 | 3 | 95.5 | 4.5 |
| 1f | E6_Reporter / Gad1_layer4 | 3 | 100 | 0 |
| 1f | E6_Reporter / Gad1_layer5 | 3 | 98.3 | 1.7 |
| 1f | E6_Reporter / Gad1_layer6 | 3 | 100 | 0 |
| 1f | E6_Reporter / PV_layer1 | 3 | 0 | 0 |
| 1f | E6_Reporter / PV_layer2/3 | 3 | 5.4 | 1.4 |
| 1f | E6_Reporter / PV_layer4 | 3 | 3.4 | 0.6 |
| 1f | E6_Reporter / PV_layer5 | 3 | 2.3 | 0.6 |
| 1f | E6_Reporter / PV_layer6 | 3 | 1.8 | 0.3 |
| 1f | E6_Reporter / SST_layer1 | 3 | 0 | 0 |
| 1f | E6_Reporter / SST_layer2/3 | 3 | 0 | 0 |
| 1f | E6_Reporter / SST_layer4 | 3 | 7.5 | 4.6 |
| 1f | E6_Reporter / SST_layer5 | 3 | 5.9 | 2.6 |
| 1f | E6_Reporter / SST_layer6 | 3 | 2.9 | 2.9 |
| 1f | E6_Reporter / VIP_layer1 | 3 | 0 | 0 |
| 1f | E6_Reporter / VIP_layer2/3 | 3 | 92.5 | 2.6 |
| 1f | E6_Reporter / VIP_layer4 | 3 | 76.3 | 8.8 |
| 1f | E6_Reporter / VIP_layer5 | 3 | 76.7 | 3.8 |
| 1f | E6_Reporter / VIP_layer6 | 3 | 96.2 | 3.8 |
| 1f | E2_Density Gad1+_layer1 | 3 | 0.3 | 0.5 |
| 1f | E2_Density Gad1+_layer2/3 | 3 | 12.5 | 1 |
| 1f | E2_Density Gad1+_layer4 | 3 | 30.9 | 3.1 |
| 1f | E2_Density Gad1+_layer5 | 3 | 35 | 1.4 |
| 1f | E2_Density Gad1+_layer6 | 3 | 18.8 | 4.8 |
| 1f | E2_Density Gad1-_layer1 | 3 | 0 | 0 |
| 1f | E2_Density Gad1-_layer2/3 | 3 | 0.7 | 0.1 |
| 1f | E2_Density Gad1-_layer4 | 3 | 0 | 0 |
| 1f | E2_Density Gad1-_layer5 | 3 | 1.1 | 0.6 |
| 1f | E2_Density Gad1-_layer6 | 3 | 0.7 | 0.9 |

| F | Quantification | n | mean | s.e.m |
|---|---|---|---|---|
| 1f | E5_Density Gad1+_layer2/3 | 3 | 22.2 | 1.5 |
| 1f | E5_Density Gad1+_layer4 | 3 | 12.2 | 0.5 |
| 1f | E5_Density Gad1+_layer5 | 3 | 18.6 | 0.3 |
| 1f | E5_Density Gad1+_layer6 | 3 | 8.4 | 1.9 |
| 1f | E5_Density Gad1-_layer1 | 3 | 0.3 | 0.4 |
| 1f | E5_Density Gad1-_layer2/3 | 3 | 0.3 | 0.4 |
| 1f | E5_Density Gad1-_layer4 | 3 | 0.6 | 0 |
| 1f | E5_Density Gad1-_layer5 | 3 | 34.7 | 3 |
| 1f | E5_Density Gad1-_layer6 | 3 | 0.6 | 0.8 |
| 1f | E6_Density Gad1+_layer1 | 3 | 0 | 0 |
| 1f | E6_Density Gad1+_layer2/3 | 3 | 34.7 | 7.2 |
| 1f | E6_Density Gad1+_layer4 | 3 | 36.8 | 1.3 |
| 1f | E6_Density Gad1+_layer5 | 3 | 13.7 | 6.6 |
| 1f | E6_Density Gad1+_layer6 | 3 | 13.6 | 1 |
| 1f | E6_Density Gad1-_layer1 | 3 | 0 | 0 |
| 1f | E6_Density Gad1-_layer2/3 | 3 | 0 | 0 |
| 1f | E6_Density Gad1-_layer4 | 3 | 0.6 | 0.9 |
| 1f | E6_Density Gad1-_layer5 | 3 | 0 | 0 |
| 1f | E6_Density Gad1-_layer6 | 3 | 0.6 | 0.9 |
| 2b | E2_Reporter / PV_layer1-6 | 4 | 91.6 | 0.9 |
| 2b | E2_Reporter / PV_layer1-6 | 8 | 90.8 | 1.1 |
| 2b | E2_PV / reporter_layer1-6 | 19 | 75.7 | 2.9 |
| 2c | E2_Reporter at .5_layer1-6 | 3 | 10 | 1.1 |
| 2c | E2_Reporter at 2_layer1-6 | 3 | 33 | 10.6 |
| 2c | E2_Reporter at 5_layer1-6 | 3 | 96.5 | 8.4 |
| 2c | E2_Reporter / PV at .5_layer1-6 | 3 | 86.4 | 0.5 |
| 2c | E2_Reporter / PV at 2_layer1-6 | 3 | 86.6 | 0.6 |
| 2c | E2_Reporter / PV at 5_layer1-6 | 3 | 91.3 | 0.8 |
| 2d | E2_Reporter / PV 1-15_layer1-6 | 4 | 56.7 | 2.3 |
| 2d | E2_Reporter / PV 7-15_layer1-6 | 5 | 67.2 | 1.5 |
| 2d | E2_Reporter / PV 10-15_layer1-6 | 3 | 81.7 | 1.1 |
| 2d | E2_Reporter / PV 4-7_layer1-6 | 2 | 60.9 | 1.9 |
| 2d | E2_Reporter / PV 7-10_layer1-6 | 5 | 78.1 | 1.8 |
| 3a | E2_Reporter / PV_layer1-6 | 4 | 85.3 | 0.6 |
| 3a | E2_PV / reporter_layer1-6 | 4 | 87.7 | 1.2 |
| 3b | E2_Delta _layer1-6 | 14 | 8.2 | 0.8 |
| 4a | E2_Rat_layer1-6 | 1 | 93 | na |
| 4a | E2_Marmoset_layer1-6 | 4 | 91.8 | 3.1 |
| 4a | E2_Macaque_layer1-6 | 4 | 87.3 | 0.5 |
| 4b | E2_Human_reporter_layerna | 44 | 72.7 | na |
| 4b | E2_Human_C1V1_layerna | 10 | na | na |
| 5a | E11_Reporter / Gad1_layer1-6 | 2 | 94.7 | 1.3 |
| 5a | E11_Reporter / PV_layer1-6 | 3 | 91.3 | 1.6 |
| 5a | E14_Reporter / Gad1_layer1-6 | 2 | 92.5 | 2.7 |
| 5a | E14_Reporter / PV_layer1-6 | 3 | 93.6 | 1.1 |
| 5a | E22_Reporter / Gad1_layer1-6 | 2 | 97.6 | 2.4 |
| 5a | E22_Reporter / PV_layer1-6 | 3 | 92.9 | 2 |
| 5a | E29_Reporter / Gad1_layer1-6 | 2 | 95.8 | 1.2 |
| 5a | E29_Reporter / PV_layer1-6 | 2 | 94.3 | 1.3 |
| 5b | E22_Reporter / Gad1_layer1-6 | 2 | 91.8 | 1.6 |
| 5b | E22_ Reporter / PV_layer1-6 | 6 | 81.3 | 1.6 |

**Supplementary Table 2.** Table containing the metadata associated with each of the quantification plot presented in this the figures of this manuscript.
